# Spike-timing-dependent plasticity can account for connectivity aftereffects of dual-site transcranial alternating current stimulation

**DOI:** 10.1101/2020.10.16.342105

**Authors:** Bettina C. Schwab, Peter König, Andreas K. Engel

**Affiliations:** Department of Neurophysiology and Pathophysiology, University Medical Center Hamburg-Eppendorf, Germany; Berlin Institute for Advanced Study, Germany; Institute of Cognitive Science, University of Osnabrück,Germany

**Keywords:** Transcranial alternating current stimulation, Spike-timing-dependent plasticity, Electroencephalogram, Functional connectivity, Entrainment

## Abstract

Transcranial alternating current stimulation (tACS), applied to two brain sites with different phase lags, has been shown to modulate stimulation-outlasting functional EEG connectivity between the targeted regions. Given the lack of knowledge on mechanisms of tACS aftereffects, it is difficult to further enhance effect sizes and reduce variability in experiments. In this computational study, we tested if spike-timing-dependent plasticity (STDP) can explain stimulation-outlasting connectivity modulation by dual-site tACS and explored the effects of tACS parameter choices. Two populations of spiking neurons were coupled with synapses subject to STDP, and results were validated via a re-analysis of EEG data. Our simulations showed stimulation-outlasting connectivity changes between in- and anti-phase tACS, dependent on both tACS frequency and synaptic conduction delays. Importantly, both a simple network entraining to a wide range of tACS frequencies as well as a more realistic network that spontaneously oscillated at alpha frequency predicted that the largest effects would occur for short conduction delays between the stimulated regions. This finding agreed with experimental EEG connectivity modulation by 10 Hz tACS, showing a clear negative correlation of tACS effects with estimated conduction delays between regions. In conclusion, STDP can explain connectivity aftereffects of dual-site tACS. However, not all combinations of tACS frequency and application sites are expected to effectively modulate connectivity via STDP. We therefore suggest using appropriate computational models and/or EEG analysis for planning and interpretation of dual-site tACS studies relying on aftereffects.

**Highlights:**

- Network model with STDP explains EEG connectivity change after dual-site tACS
- Effects are predicted to depend on tACS frequency and conduction delays
- EEG data confirm dependence on conduction delays between regions
- Model can be used to estimate and maximize experimental effects

## 1 Introduction

Transcranial alternating current stimulation (tACS) became a popular tool to modulate both oscillations and oscillatory connectivity in the brain as measured by different imaging modalities [1–3]. As the measurement of extracellular potentials during tACS, including EEG and MEG, is hampered by large and hard-to-predict stimulation artifacts [4, 5], measurement of stimulation-outlasting aftereffects turned into the standard for EEG and MEG [6, 7]. So far, mechanisms mediating these aftereffects are not identified yet, but spike-timing-dependent plasticity (STDP) has been proposed as a candidate [8]. For EEG aftereffects in alpha- and beta-power, experimental evidence supports the involvement of STDP [9, 10].

Applying electric fields with different phase lags to two different regions of the brain (dual-site tACS) seems ideal to exploit the plasticity of synapses between the targeted regions to induce aftereffects. Indeed, we [6] recently found that tACS at 10 Hz, applied in a focal montage to each hemisphere, transiently modulates stimulation-outlasting functional EEG connectivity in the alpha range depending on the tACS phase lag between the two applied electric fields. Specifically, functional connectivity increased after in-phase stimulation (tACS phase lag 0) compared with anti-phase stimulation (tACS phase lag *π*).

Here, we aim to test in small computational models whether STDP can account for the experimentally observed connectivity change after dual-site 10 Hz tACS [6]. We thereby assume that tACS entrains at least a subset of the targeted neurons, as shown to be the case in macaques [11, 12]. First, we use a simplified model susceptible to many tACS frequencies to investigate which experimental settings maximize the size of effects in a given direction. Second, we use a more realistic network model to directly validate the outcome of 10 Hz stimulation with existing EEG data [6] (Fig. 1).

**Figure 1:**
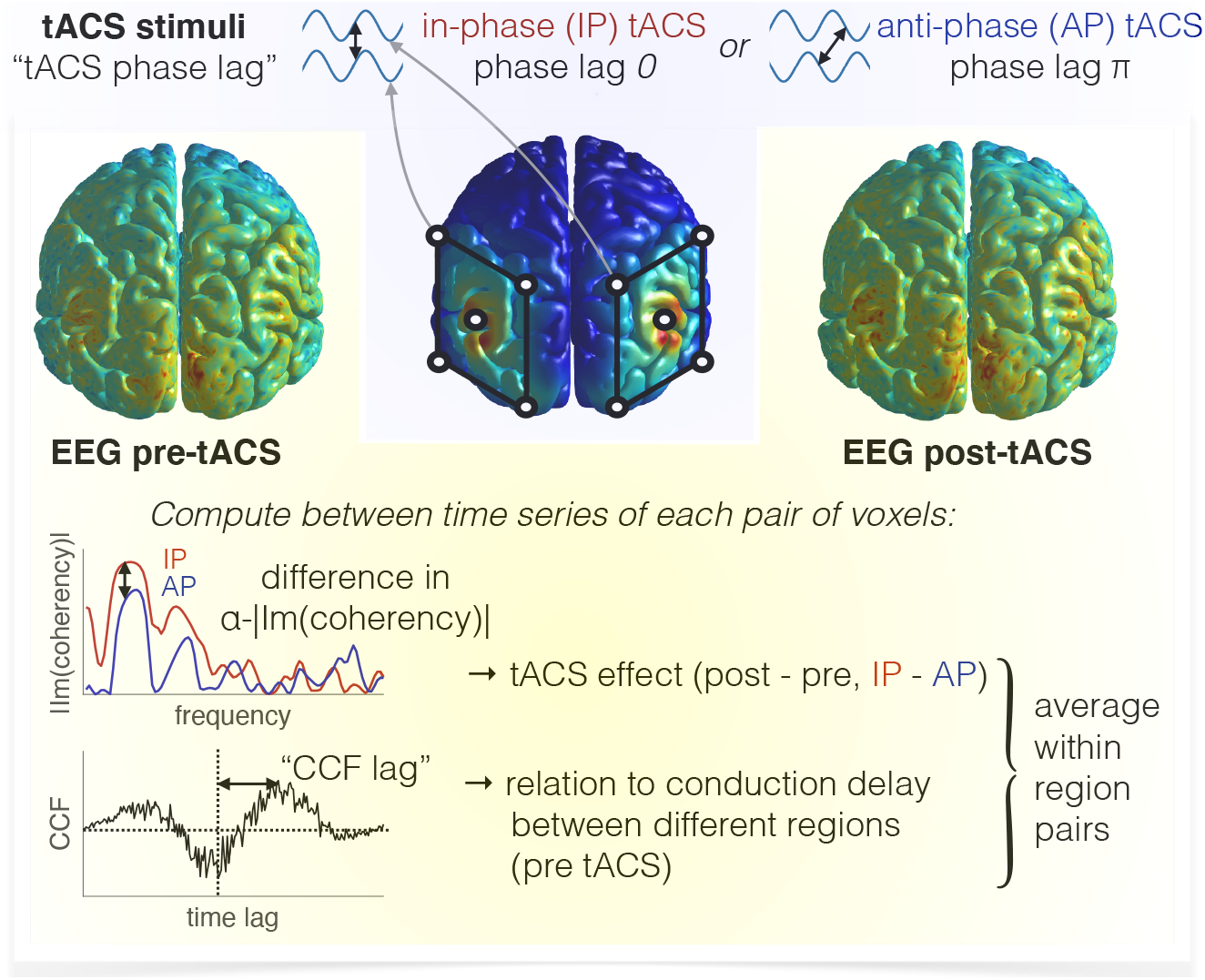
Summary of experimental data for validation of the model. tACS was applied at 10 Hz in- (tACS phase lag 0) and anti-phase (tACS phase lag *π*), while EEG was recorded before and after each stimulation. The difference in imaginary coherence in the alpha-band between in- and anti-phase stimulation, each computed as the difference between post- and pre-tACS, was considered as the main tACS effect. Pre-tACS data served to estimate conduction delays between regions by detecting the peak absolute value of each cross-correlation function (CCF) and taking the absolute value of its time lag.

## 2 Materials and Methods

We used two small network models of spiking neurons to test the effect of different dual-site tACS protocols on connectivity changes due to STDP. Each network consisted of two populations with 1000 Izhikevich neurons each [13] that followed the ordinary differential equations

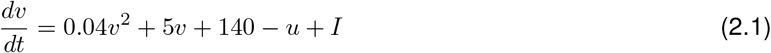

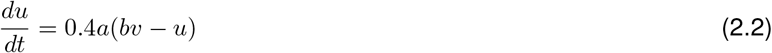

with the auxiliary after-spike resetting

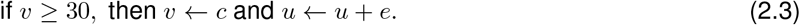

Although the models are dimensionless, the variables *v* (transmembrane voltage) and *t* (time) can be interpreted in units of mV and ms, respectively. The membrane recovery variable *u* provides negative feedback to *v*. After *v* reaches the apex of a spike (30 mV), both *v* and *u* are reset. The parameters *a*, *b*, *c*, and *e* describe the time scale of *u*, the sensitivity of *u* to subthreshold fluctuations of *v*, the after-spike reset value of *v*, and the after-spike reset of *u*, respectively, which differ for neuron types. *I* represents input currents to the neuron, summing up the synaptic input from other neurons, random thalamic input, and tACS currents.

Depending on the network’s architecture, neurons were synaptically coupled with fixed axonal conduction delays. After a spike in a presynaptic neuron, each postsynaptic neuron received an input of size *w* with delay *d*. Excitatory synapses between the two populations were subject to STDP. Connections between the populations were set to *w* = 0.01 in order to preserve the dynamics of each population but weakly couple the populations. Synaptic weight change Δ*w* followed the rule

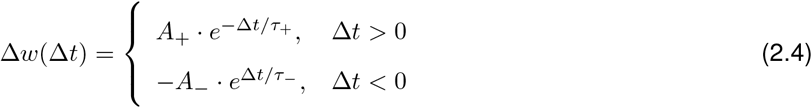

with the difference between post- and presynaptic spike timing at the synapse Δ*t = t_post_ − t_pre_*, the maximum potentiation *A*_+_ = 0.015, the maximum depression *A*_−_ = 0.007, the time constant for potentiation *τ*_+_ = 13*ms*, and the time constant for depression *τ*_−_ = 34*ms*. This rule was found experimentally by Froemke and Dan [14] in slices of rat visual cortex, layer 2/3, and is shown in Fig. 2A. Characteristics of this rule — in particular, *A*_+_ > *A*_−_ and *τ*_+_ < *τ*_−_ — have been corroborated by many other studies [15–17].

**Figure 2:**
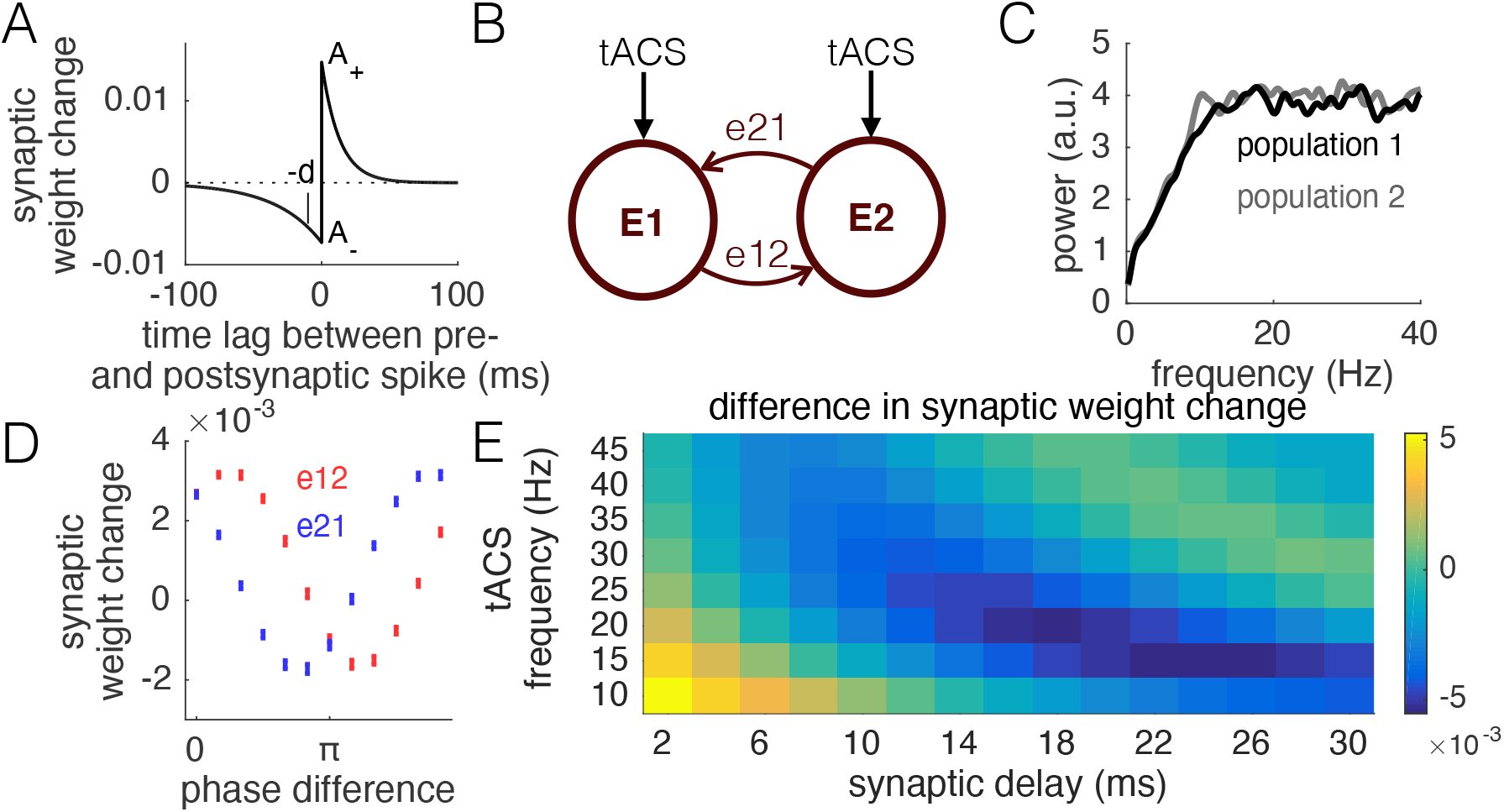
A simple model of two spiking neuron populations (Network 1). A: STDP rule based on [14]. The lag Δ*t* between pre- and postsynaptic spike time *t_pre_* and *t_post_* was defined as Δ*t* = *t_post_* − *t_pre_*. Thus, presynaptic spikes occurring before postsynaptic spikes led to potentiation of synaptic weights. B: Network 1 consisted of two populations with coupling only to neurons of the other population. C: Without stimulation, the network did not oscillate at a particular frequency. D: Stimulation of the two populations at different tACS phase lags (defined as the phase of tACS to population 2 minus the phase of tACS to population 1) affected connectivity (mean ± STD) differently from population 1 to population 2 (*e*12) and vice versa (*e*21), shown here for *d* = 10 *ms*. For example, if population 1 was stimulated 10 *ms* or a phase difference of *π/*5 before population 2, *e*12 was optimally potentiated. At phase 0 (“in-phase tACS”) and phase *π* (“anti-phase tACS”), the system is symmetric, thus *e*12 and *e*21 showed comparable changes in synaptic weights. E: The average difference in synaptic weight change between in- and anti-phase stimulation depended on both tACS frequency and synaptic delay of connections *e*12 and *e*21. For low tACS frequencies and short delays, in-phase stimulation increased synaptic weights compared with anti-phase stimulation.

Importantly, synaptic weights will on average decrease in the case of independent Poisson firing 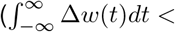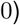. In contrast, if Δ*t* is small (say, distributed equally between *−ε* and *ε* with small, positive *ε*),average synaptic weights increase 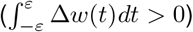.Since a presynaptic spike at time *t*_1_ arrives at the synapse at time *t*_1_ + *d*, with the synaptic conduction delay *d* of this synapse, sharp synchrony of somatic pre- and postsynaptic spikes at time *t*_1_ leads to Δ*t = t_post_ − t_pre_ = t*_1_ *−* (*t*_1_ + *d*) = *−d*, and is expected to depress synapses (see Fig. 2A). However, spike synchrony with sufficient jitter may lead to a net potentiation (e.g., 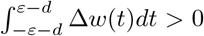 for sufficiently small *d* and sufficiently large *ε*). For details on this mechanism, see [18].

### Network 1

First, we implemented a network without connections within each population (Fig. 2B), producing random firing in the absence of tACS, and tACS-phase locked activity in the presence of tACS at various frequencies (Supplement A). All neurons were assigned *b* = 0.2, *c* = −65*mV* , and *e* = 8.The parameter *a* differed between neurons and was randomly set for each neuron following a Gaussian distribution *A* with mean 0.04 and standard deviation (STD) 0.015. Negative values of *a* were set to their absolute value. Every millisecond, 40 neurons of each population were chosen randomly to receive an input of 20*a/Ā*. Thus, the larger the parameter *a* of a neuron, the larger its thalamic input. This combination of the parameter *a* and thalamic input led to a flat power spectrum (Fig. 2C) and a realistic distribution of firing rates (Supplement A). Each neuron was assigned 100 excitatory connections with low weight *w* = 0.01 to random neurons of the other population with delay *d*. tACS was modeled as sinusoidal current input with amplitude 2 to all neurons, with tACS phase lag 0 or *π* between the populations. The tACS frequency was modulated between 10 and 45 Hz. Network 1 was simulated for two seconds with tACS, while synaptic weights of the last second were subject to analysis, in which the network was in a steady state. Synaptic weights were compared before and after this time segment, representing the difference in structural connectivity before and after stimulation.

### Network 2

Next, we adapted the network to a more realistic architecture that produces spontaneous alpha oscillations based on Izhikevich [19] (Fig. 3A,C). Each population was divided into an excitatory subpopulation of 800 neurons and an inhibitory subpopulation of 200 neurons. Excitatory neurons were assigned *b* = 0.2, *c* = −65*mV* , *e* = 8, and *a* drawn randomly from a Gaussian distribution with mean 0.04 and STD 0.01, where negative values were set to their absolute value. Parameters of all inhibitory neurons were set to *a* = 0.2, *b* = 0.2, *c* = −65*mV* , and *e* = 2. Connectivity within each population was similar to what was chosen in Izhikevich [19]: each excitatory neuron projected to 60 random neurons, which could be excitatory or inhibitory, with fixed weight *w* = 6. Delays for those connections were distributed uniformly between 1 ms and 10 ms. Inhibitory neurons connected to 100 random excitatory neurons with fixed weight *w* = −5 and delay *d* = 1*ms*. This choice of connectivity and parameters led to realistic distributions of firing rates (Supplement B) and spontaneous emergence of alpha oscillations (Fig. 3B,D). Each excitatory neuron additionally connected to 100 random excitatory neurons of the other population with low weight *w* = 0.01 (Fig. 3A). All synapses between the populations were subject to STDP. Each millisecond, 30 random neurons of each population received a thalamic input of size 30. Since Network 2 easily entrained to 10 Hz tACS, related to its intrinsic oscillation, it was stimulated at 10 Hz with a sinusoidal current of amplitude 1 to all excitatory neurons. Functional connectivity was simulated as the alpha-coherence (8 – 12 Hz) between the average transmembrane voltage of each population and its difference between in- and anti-phase tACS was computed. Network 2 was simulated for four seconds with tACS and aftereffects of three seconds with the latter being subject to analysis of synaptic weights and stimulation-outlasting effects on functional connectivity. The simulation of neural activity after stimulation offset is here necessary to compute functional connectivity, which — in contrast to structural connectivity — requires time series of activity. Also connectivity during the last three seconds of stimulation may be analyzed, but is not the focus of this article. Comparable to Network 1, values of synaptic weights were evaluated after discarding the first second, between the first time step of the second second and the last time step of stimulation, representing structural connectivity change with stimulation.

**Figure 3:**
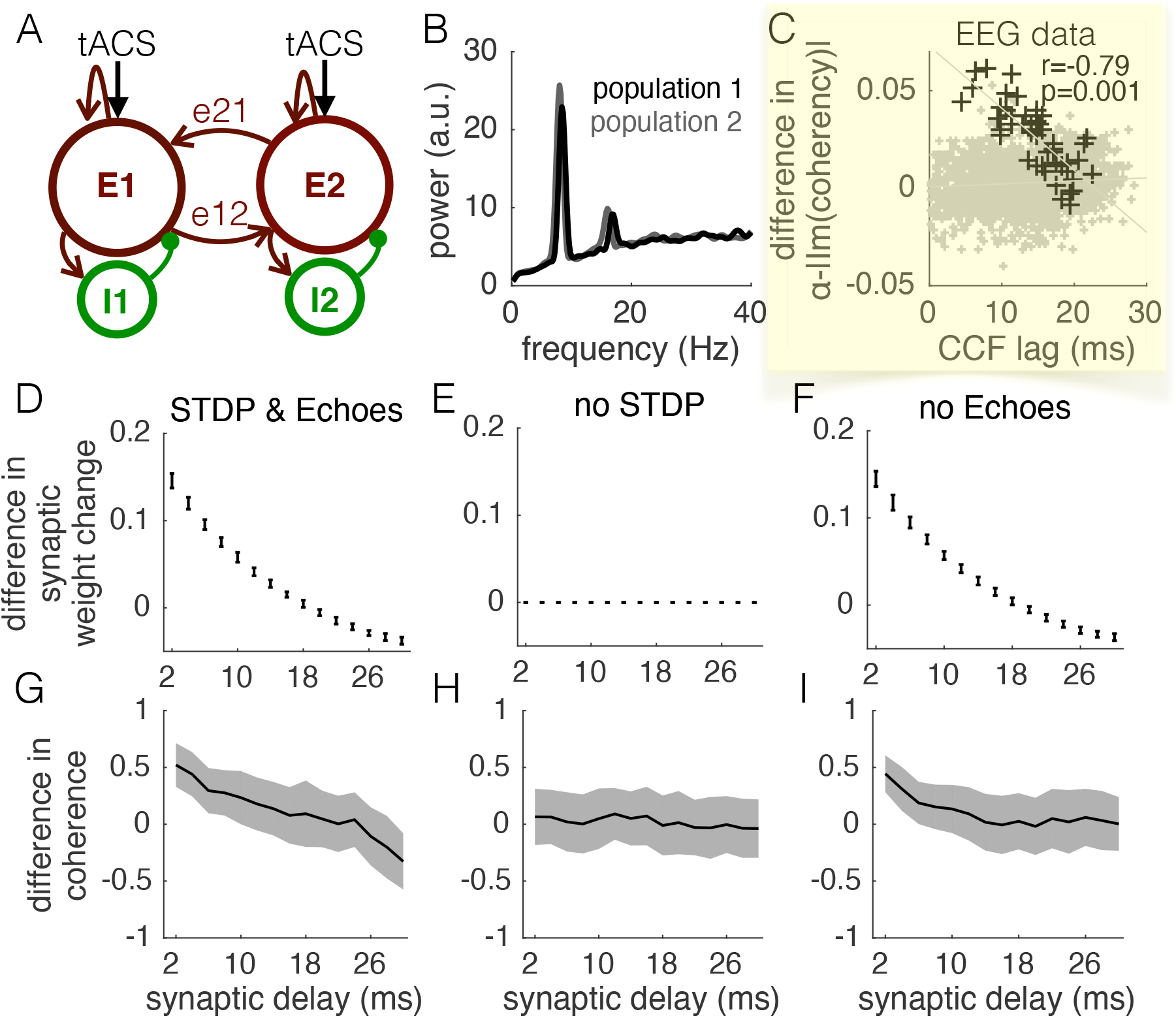
Network 2, oscillating at alpha frequency due to realistic connectivity, confirmed the results and was in agreement with experimental data. A: Architecture of Network 2. Each population consisted of an excitatory and an inhibitory subpopulation. Excitatory neurons coupled also between the two populations, and these inter-population synapses were subject to the STDP rule shown in Fig.2A. B: Network 2 showed spontaneous alpha oscillations due to intrinsic coupling within each population. C: Experimental connectivity modulation dependent on the estimated conduction delay (CCF time lag). All possible pairs of ROIs are shown as small gray crosses. Stimulated pairs (black) showed a negative relation between connectivity modulation and CCF time lag. D-F: Difference in average synaptic weight change (mean ± STD) of synapses between the populations between in- and anti-phase stimulation for the system with both STDP and entrainment echoes (D), with only entrainment echoes (E) and with only STDP (F). G-I: Respective differences in functional connectivity (mean ± STD) between simulated average transmembrane voltages of in-and anti-phase stimulation.

All simulations were run in MATLAB (The MathWorks Inc). The delay of synapses between the two populations was parametrically modulated between 2 and 30 ms. Inter-spike intervals followed a power law tail distribution as seen experimentally in single neuron recordings [20] (Supplement C). To test for the effects of STDP, the learning rule was set to zero weight change for all time lags. To remove entrainment echoes, all transmembrane voltages were reset to their original values after tACS-offset.

### Experimental validation

We also compared the simulations with our experimental data. Seven regions of interest (ROIs) in occipito-parietal areas of each hemisphere were experimentally targeted by tACS, defined as ROIs with average differences in vectorial field strength between in- and anti-phase tACS above 0.08 *V/m* [6]. Thus, we were able to investigate 49 interhemispheric connections between stimulated ROIs, possibly differing in their conduction delays. EEG data were projected to source space using ELORETA and time series in each voxel were extracted as described in [6]. For all voxel pairs, we computed the absolute value of the imaginary coherence in between 9 and 11 Hz (*α* − | Im(*coherency*)|), and subtracted values between post- and pre-tACS, as well as between in- and anti-phase tACS equivalently to [6]. Additionally, for all voxel pairs of EEG data pre-tACS, we computed the cross-correlation function (CCF), determined its absolute value, and extracted the absolute value of the time lag |*τ*| of its maximum (see also Fig. 1): |*CCF*(*τ*)| = max(|*CCF* (*t*)|). Both *α* − | Im(*coherency*)| and |*τ*| (“CCF lag”) were averaged for pairs between the same ROIs, and the averages were set in relation using Pearson correlation. Permutation statistics were used to compute the *p*-value for the correlation (Supplement D).

### Ethics Statement

The ethics committee of the Medical Association Hamburg approved the previous experimental study [6] which served as the data set for experimental validation here.

### Data and Code Availability Statement

Code to simulate Network 1 and 2 as well as EEG connectivity data for validation can be requested by e-mail from the corresponding author.

## 3 Results

We first simulated two populations of randomly spiking neurons with weak connections subject to STDP (Network 1, Fig. 2), which entrained to any tACS frequency in the range 10 *–* 45 Hz (Supplement A). Synapses between the populations were subject to STDP (Fig. 2A), and both populations were stimulated by external tACS currents (Fig. 2B). Average transmembrane voltages did not exhibit a strong oscillation, but approximately constant power between 10 and 45 Hz (Fig. 2C). 10 Hz tACS at different tACS phase lags led to direction-specific changes in mean synaptic weights (Fig. 2D). As expected, weight changes were symmetric only for tACS phase lags 0 (“in-phase stimulation”) and *π* (“anti-phase stimulation”). We focused on these two special cases and, from then on, on average synaptic weights for all connections of *e*12 and *e*21. In particular, we were interested in the difference in average synaptic weight change between in- and anti-phase stimulation, which is shown in Fig. 2E for different combinations of stimulation frequency and synaptic delay. Clear effects in the positive direction direction — in-phase stimulation leads to an increase in synaptic weight compared with anti-phase stimulation — were found for low frequencies and short delays. With increasing frequency and delay, the effects diminished and eventually reversed.

Do our results depend on a realistic connectivity architecture and the presence of intrinsic oscillations, as for example found in the alpha-band for resting-state EEG? To answer this question, we extended the network with inhibitory subpopulations as well as intrinsic coupling (Network 2, Fig. 3A), which led to strong spontaneous alpha oscillations (Fig. 3B). Network 2 easily entrained to tACS at 10 Hz (Supplement B), providing an environment comparable to our previous experimental setup [6]. Re-analysis of our experimental data revealed that the difference in *α* − | Im(*coherency*)| between in- and anti-phase stimulation, the main tACS effect, dropped with the conduction delay between ROIs as estimated by CCF lags before application of tACS (Fig. 3C; for details, see Supplement E). In fact, our model predicted the same finding: differences in both simulated structural connectivity (Fig. 3D) and simulated functional connectivity (Fig. 3G) decreased with the conduction delay. At delay (20.2±5.2) ms, the change in functional connectivity in our model was 0, in accordance with the experimental data ((22.8±7) ms). Also Network 1 is qualitatively in agreement with this experimental data: For 10 Hz stimulation (Fig. 2E), the difference in synaptic weights decreased monotonically with increasing synaptic delays. Additional modulation of the tACS frequency in Network 2 is shown in Supplement F. Finally, we tested in Network 2 whether the change in simulated functional connectivity actually related to the simulated change in synaptic weights due to STDP, or was rather a result of entrainment echos. Removal of STDP (Fig. 3E,H) led to an absence of effects, while removal of entrainment echoes (Fig. 3F,I) only attenuated the effects seen with both STDP and entrainment echoes (Fig. 3D,G). Therefore, in this model, STDP seems necessary to induce the experimentally observed effects, while entrainment echoes can further enhance effect sizes.

## 4 Discussion

With small network models of spiking neurons, we demonstrated that aftereffects of dual-site tACS on functional connectivity at an intrinsic frequency of the network can be explained by STDP of synapses between the stimulated ROIs, possibly in combination with entrainment echoes. Our work thereby adds to the existing literature suggesting plasticity to underly aftereffects of single-site tACS [8–10] and provides a framework to test effects of stimulation parameters on stimulation-outlasting connectivity modulation. Critically, our first model, easily entraining to frequencies between 10 and 40 Hz, further suggests that the effects in synaptic weight change depend on tACS frequency and synaptic delays, with the largest effects for low tACS frequencies and short delays. Here, we assume that synaptic weight changes during stimulation induce changes in functional connectivity that can be measured after stimulation. Our second model, oscillating spontaneously at alpha frequency, showed that adding biological detail to the network left the negative relation of connectivity modulations with synaptic delays at 10 Hz stimulation intact. Functional connectivity, computed in time windows of several seconds after stimulation, followed the direction of structural connectivity change and could then directly be compared to our existing EEG data [6]. Re-analysis of the data confirmed the dependence of functional and structural connectivity modulation on the estimated conduction delay as seen in both computational models, and thereby explained the heterogeneity of experimental results across ROI pairs.

Indeed, most robust aftereffects of dual-site tACS on functional connectivity described in the literature have been found for theta frequencies [21]. Literature on aftereffects of single-site tACS on EEG power (e.g., [8–10]) has so far focused on the alpha- and beta-band. In contrast, neural effects during tACS may be directly related to entrainment of spiking [11, 12] rather than plasticity of synapses and could therefore depend on factors different from synaptic delays and stimulation frequencies. The mentioned studies [11, 12] found comparable entrainment effects across different tACS frequencies, including the gamma-range, which is in line with our reasoning. While extensive literature exists on the roles of conduction delays for long-range synchronization [22–24], their roles for dual-site tACS have — to our knowledge — so far not been analyzed systematically.

It is not the aim of this study to present a dynamic model with realistic cortical dynamics and effect sizes of tACS. Instead, the model relies on the assumption that spike *timing* is relevant for connectivity modulation. Thus, firing rate distributions as well as the STDP rule are critical and have to be adjusted to experimental data. In particular, we simulated only short time intervals to detect the direction of connectivity change under stimulation. Moreover, depending on factors like montage, amplitude, or brain state, tACS is expected to entrain only a subset of the targeted neurons. We simplify the situation by choosing tACS amplitudes large enough to entrain the majority of cells (Supplement A, B). Nevertheless, this discrepancy does not affect our conclusions, as the described mechanism of connectivity modulation holds for the subset of neurons that are entrained, and this subset may drive the experimentally observed changes in functional connectivity.

## 5 Conclusions

Our computational models demonstrate that STDP can explain experimentally observed EEG connectivity changes after dual-site stimulation at the network’s intrinsic frequency. The direction of aftereffects is expected to depend on both tACS frequency and conduction delay between the targeted ROIs. Computational modeling and EEG connectivity analysis to estimate conduction delays may in general be used to predict the sign of expected connectivity modulation by tACS to decrease variability and maximize effect sizes in future experimental studies — factors that are among the main challenges for tACS research [3].

## Acknowledgement

This work has been supported by DFG, SFB 936/project A3, and the Berlin Institute for Advanced Study. We thank Marina Fiene for helpful discussions as well as Jan-Ole Radecke and Marina Fiene for proof-reading of the manuscript.

## Supplement

### Supplement A

**Supplement A:**
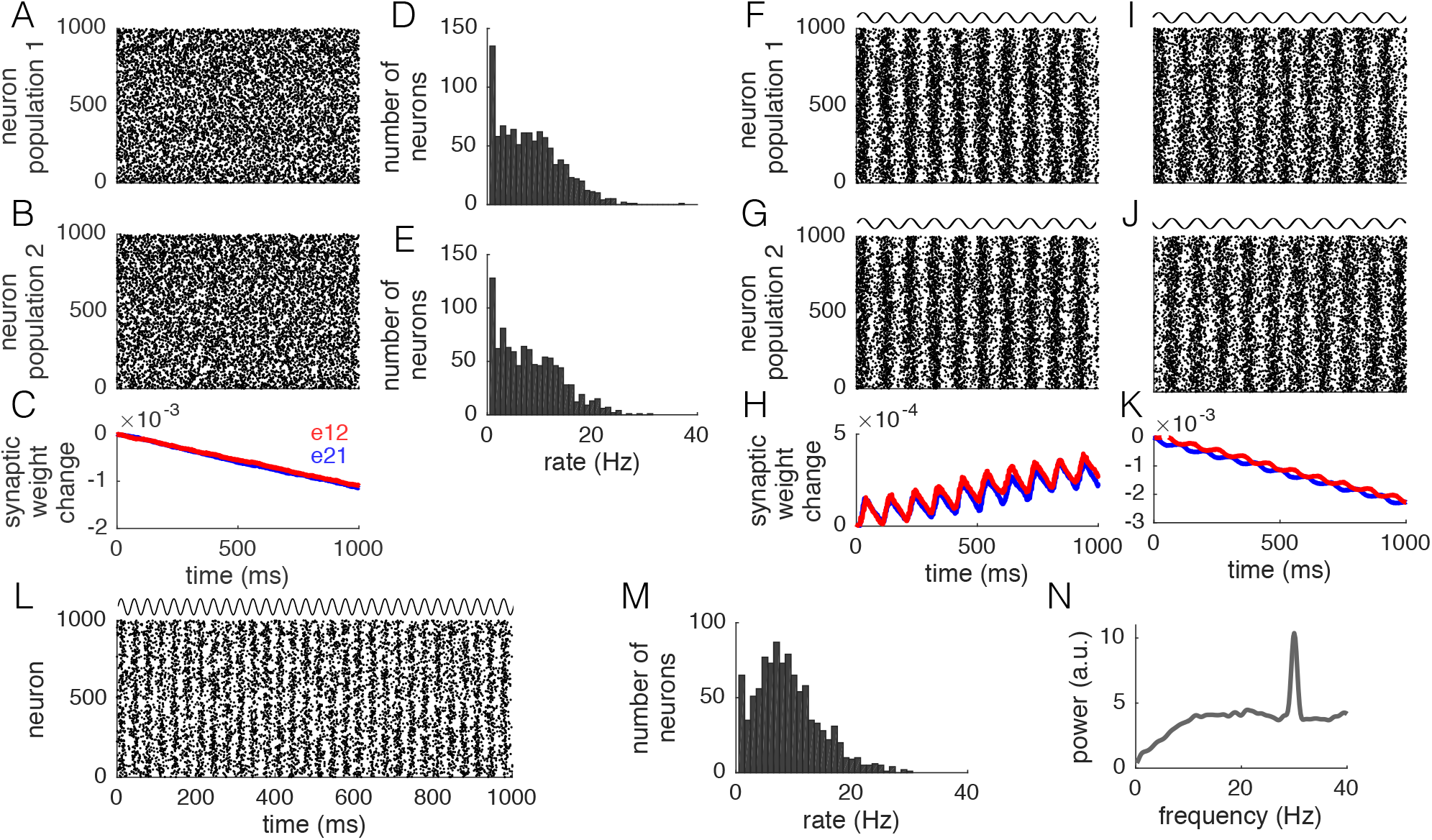
Examples for Network 1 with *d* = 2 *ms.* A, B: Firing patterns of neurons in population 1 and 2, respectively. No prominent oscillation is visible. C: Average synaptic weights of the connections between the two randomly spiking populations decreased. D, E: Firing rates of both populations were distributed in a wide range between 0 and 40 Hz. F, G: In-phase tACS led to synchronous entrainment of the two population and a net potentiation of synaptic weights (H). I, J: In contrast, anti-phase stimulation entrained the two populations at tACS phase lag *π* and induced a net depression of synaptic weights (K). L: Also tACS at higher frequencies entrained the network, here shown for 30 Hz. While firing rates were still distributed mainly around 10 Hz (M), average transmembrane voltages showed a power peak at 30 Hz (N).

### Supplement B

**Supplement B:**
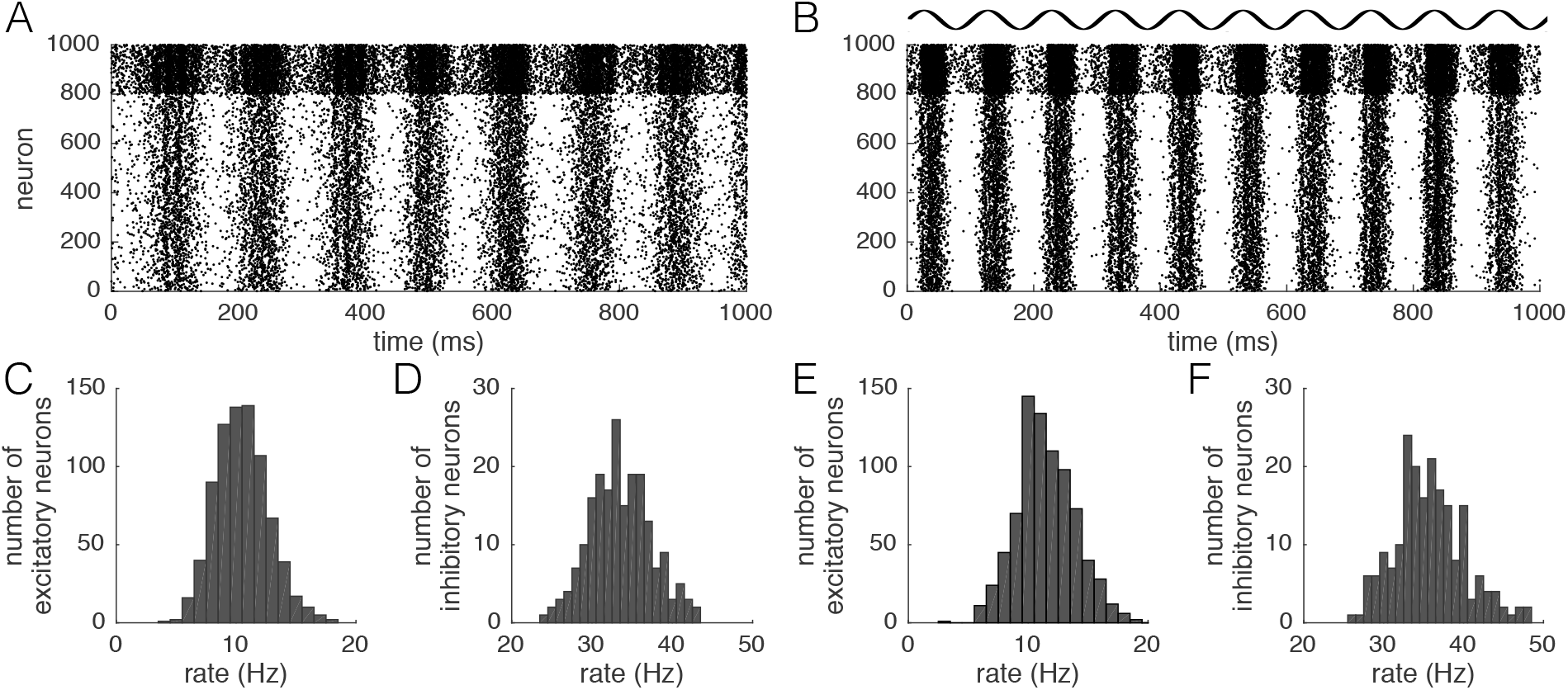
Entrainment in Network 2. A: Without tACS, the network spontaneously oscillated at around 8 Hz. B: 10 Hz tACS entrained the network to the stimulated frequency and phase. Firing rate distributions for excitatory (C, E) and inhibitory (D, F) neurons are comparable between the un-stimulated (C, D) and the stimulated (E, F) network.

### Supplement C

#### Power-law distributions of inter-spike intervals

Inter-spike intervals (ISIs) of simulated neurons showed similar power law distributions as experimentally observed ISIs [20]. The power law exponent of the distribution’s tail (inter-spike intervals ≥ 60 ms) was −2.1 ±0.1 (mean ± STD) and −3.8±1.1 for Network 1 and 2, respectively.

**Supplement C:**
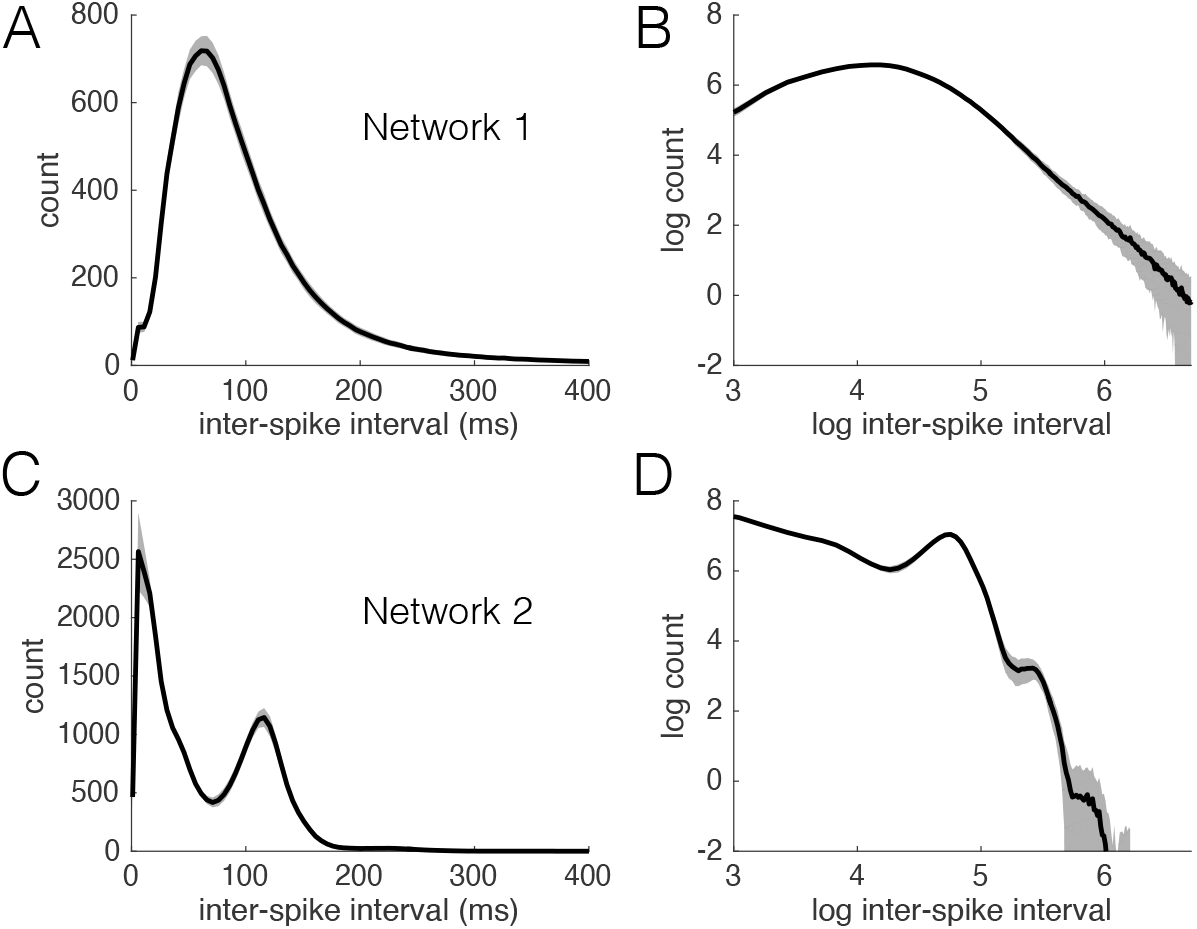
ISIs of simulated neurons (A, B: Network 1; C, D: Network 2) followed a power law distribution for large values of ISIs (mean ± STD). A, C: Distribution of ISIs, B, D: shown as log-log plot.

### Supplement D

#### Permutation statistics of Pearson correlations

We used permutation statistics to determine the *p*-value of the correlation between experimentally observed tACS effects and the estimated conduction delay (CCF lag) between the targeted regions (Fig. 3C, Supplement E). Each point in the scatter plot represents the connection between one ROI in the left hemisphere and one ROI in the right hemisphere. As ROIs contribute not only to one, but to multiple connections, values for both *α* − | lm(*coherency*)| change and CCF lag are dependent on values of other connections. We therefore did not permute connections, but the seven stimulated ROIs, keeping the dependence between connections due to shared ROIs. The seven ROIs can be permuted in 7! = 5040 different sequences. For each sequence, we computed the Pearson correlation value (“permutation *r*”) equivalently to the Pearson correlation value obtained from the unshuffled data (“test *r*”). The *p*-value could then be determined by computing the percentile of the test *r* in the distribution of permutation *r*.

### Supplement E

#### Details and robustness of experimental results

To validate our model, we re-analyzed EEG data recorded in healthy participants before and after tACS [6]. Conduction delays between ROIs were approximated by the absolute time lag of the absolute CCF peak. The differences in alpha-imaginary coherence between in” and anti-phase stimulation negatively correlated with the estimated conduction delay (*r* = −0.79, *p* = 0.001) when restricting the analysis to the stimulated pairs (Fig. 3C, black crosses). We additionally analyzed the change in absolute coherence between ROIs that were not dominated by volume conduction. Domination of volume conduction was defined for ROI pairs with CCFs that were largest at absolute time lags below 1 *ms*. Again, for stimulated pairs, condition differences in absolute coherence negatively correlated with the estimated conduction delay (*r* = −0.38, *p* = 0.03), although results were much more variable, possibly due to remaining effects of volume conduction. Finally, we considered an alternative measure for conduction delays between ROIs, the width of the largest CCF peak at half maximum. Similar to the previous measure (CCF lag), we found a negative correlation between this CCF width and differences in alpha-imaginary coherence (*r* = −0.51, *p* = 0.01), further stressing the robustness of our findings.

Furthermore, we investigated if the negative relation between CCF lags and differences in functional alpha-coupling between in- and anti-phase stimulation may relate to intrinsic factors of our data. In particular, CCF lags may be dependent on baseline levels of *α* − |lm(*coherency*)|, which in addition may correlate with the observed differences in *α* − |lm(*coherency*)|. Indeed, we found both of these correlations (panels A, B). However, the partial correlation between CCF lags and *α* − |lm(*coherency*)| differences, correcting for the influence of baseline *α* − |lm(*coherency*)|, was still strong and significant (*r* = −0.74, *p* = 0.008) when restricted to the stimulated pairs (panel C), but absent for the set of all possible pairs (*r* = −0.018, *p* = 0.29). In sum, baseline *α* − | lm(*coherency*)| may have slightly influenced the relationship between CCF lag and *α* − | lm(*coherency*)| changes, but did not solely drive the observed negative relation.

**Supplement E:**
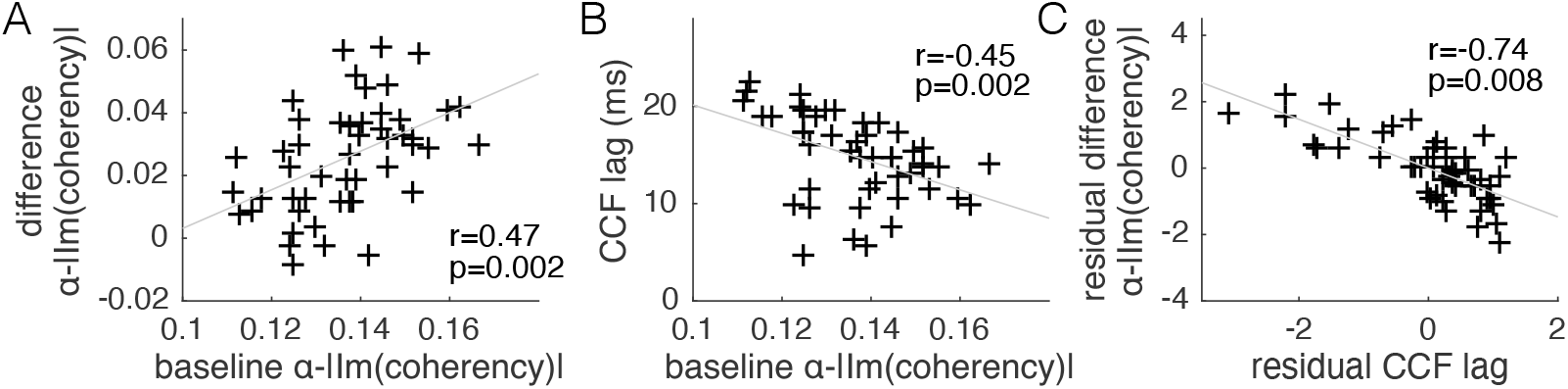
Control analyses for EEG data (stimulated pairs only). A: Scatter plot of baseline *α* − | lm(*coherency*)| and differences in *α* − |lm(*coherency*)| between in- and anti-phase stimulation. For stimulated pairs, a significant positive correlation exists. B: Scatter plot of CCF lags and baseline *α* − | lm(*coherency*)|. Stimulated pairs show a slight negative correlation. C: When correcting for the influence of baseline *α* − | lm(*coherency*)|, the negative correlation between CCF lag and *α* − | lm(*coherency*)| differences is still significant.

### Supplement F

#### Modulation of tACS frequency in Network 2 and frequency specificity of effects

In contrast to Network 1, Network 2 did not easily entrain to a wide range of tACS frequencies, impeding a direct investigation of tACS frequency dependent effects. With a tACS amplitude of 1, thus, the difference in synaptic weight change between in- and anti-phase tACS dropped with increasing synaptic delay only for tACS frequencies in the alpha range (panel A). Entraining Network 2 with a larger tACS amplitude of 10, in contrast, showed qualitatively the same dependency of synaptic weight change on tACS frequency as Network 1 (panel B), with largest effects being observed for low tACS frequencies and short synaptic delays. Finally, we also investigated the frequency-specificity of the effects of 10 Hz tACS (panel C). For each frequency, we computed a linear regression between synaptic delay and synaptic weight change, and extracted the slope of the fit. This slope was negative for all frequencies and showed a minimum at alpha frequencies, demonstrating largest tACS effects in the alpha-band.

**Supplement F:**
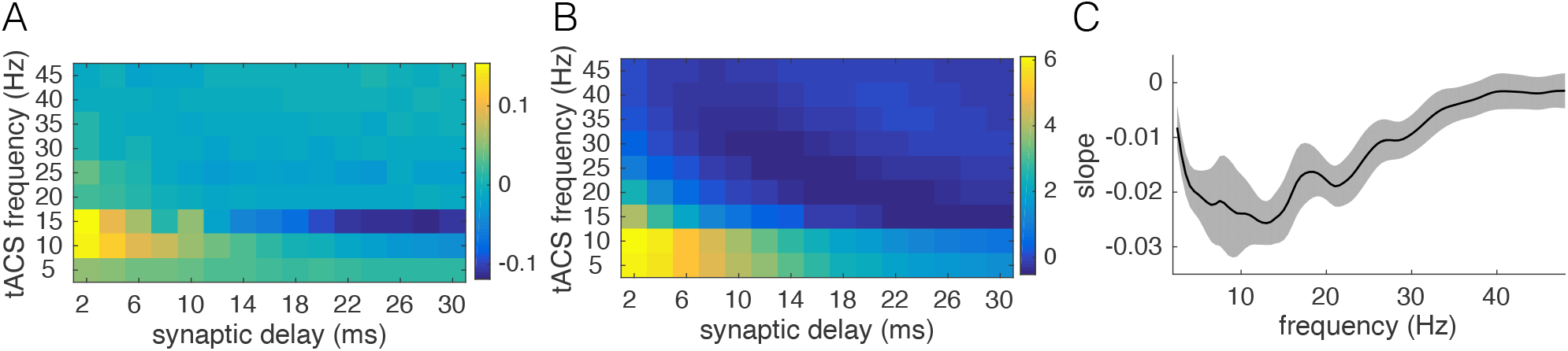
Modulation of tACS frequency in Network 2 and frequency specificity of effects. A: With the small tACS amplitude of 1 as used in Fig. 3, only alpha frequencies entrained the network and induced a tACS effect. B: With a very large tACS amplitude of 10, the network could be forced to entrain to any frequency between 5 and 45 Hz, and showed a similar dependency on tACS frequency and synaptic delay as Network 1 (see Fig. 2). C: Dependency between synaptic delay and synaptic weight change under 10 Hz tACS was largest in the alpha frequency range.

